# Widespread sex ratio polymorphism in *Caenorhabditis* nematodes

**DOI:** 10.1101/2021.10.26.465869

**Authors:** Yun Huang, Yun-Hua Lo, Jung-Chen Hsu, Tho Son Le, Fang-Jung Yang, Tiffany Chang, Christian Braendle, John Wang

## Abstract

Although equal sex ratio is ubiquitous and represents an equilibrium in evolutionary theory, biased sex ratios are predicted for certain local conditions. Cases of sex ratio bias have been mostly reported for single species, but little is known about its evolution above the species level. Here, we surveyed progeny sex ratios in 23 species of the nematode genus *Caenorhabditis*, including 19 for which we tested multiple strains. For the species with multiple strains, 5 species had female-biased and 2 had non-biased sex ratios in all strains, respectively. The other 12 species showed polymorphic sex ratios across strains. Female-biased sex ratios could be due to sperm competition whereby X-bearing sperm outcompete nullo-X sperm during fertilization. In this model, when sperm are limited allowing all sperm to be used, sex ratios are expected to be equal. However, in assays limiting mating to a few hours, most strains showed similarly biased sex ratios compared to unlimited mating experiments, except that one *C. becei* strain showed significantly reduced female bias compared to unlimited mating. Our study shows frequent polymorphism in sex ratios within *Caenorhabditis* species and that sperm competition alone cannot explain the sex ratio bias.

## Introduction

Sex ratio theory is at the core of evolutionary biology. First proposed by Charles Darwin [1, 2], and later formulated by Ronald Fisher [3], sex ratio theory posits that in a large, random mating population, when the sex ratio departs from equality, the rarer sex would have better mating prospects. Thus, genetic elements that favor the rarer sex would be favored by natural selection. Consequently, equal sex ratio should be restored and represents an evolutionary stable strategy. However, selection acts on sex ratios on different levels. Optimal individual sex ratios may differ from the population optima, as many organisms have unique life histories, social structures and parental behaviors that do not meet Fisher’s assumptions [4–6]. Biased sex ratios exist and are intriguing examples to understand selection on sex ratio.

Theories of “extraordinary sex ratios” have been proposed to explain unequal sex ratios that fit local or individual interests. Hamilton’s theory of local mate competition (LMC) predicts that in cases of highly structured populations and local mating, sex ratios would be biased toward females so that just enough males are produced to inseminate their female siblings [4]. Similarly, theories predict that sex ratio bias may arise due to local resource competition, where the sex that only consumes local resources is reduced [7]. In the case of cooperative breeding, sex ratio would be biased toward the helping sex (local resource enhancement, LRE) [8, 9]. Moreover, Trivers and Willard proposed that females in good condition or of high social ranking would produce more male offspring as their sons would inherit their advantages and have above-average breeding success [10]. These theories have successfully explained many empirical findings of sex ratio bias.

Examples of sex ratio bias across the unicellular and metazoan world are plentiful. For instance, female-biased sex ratios are commonly found in Apicomplexan parasites such as *Plasmodium malariae* [11–13] and *Toxoplasma gondii* [14], as well as in intestinal parasitic nematodes of *Heligmosomidae* [15, 16]. In Hymenoptera insects, female-biased sex ratios are found and/or tested in fig wasps [17], parasitoid wasps of *Nasonia vitripennis* [18, 19], and the *Bethylidae* family [20]. These female-biased sex ratios are good examples of LMC as these parasites are confined within their hosts. In birds, the Seychelles warbler shows facultative sex ratio bias in their offspring as they adjust production of helpers in relation to the quality of their territories [21, 22]. Male-biased sex ratios were found in African wild dogs because male helpers contribute to raising pups in the dens and increase pup survivorship [23]. These are two typical examples of LRE. Female-biased sex ratios have also been observed in red deer and wild spider monkeys with subordinate females producing more daughters but high-ranking females producing more sons [24, 25], which fits the Trivers-Willard hypothesis.

Despite the mature theories for sex ratio bias and plenty of empirical examples, our knowledge about the prevalence of sex ratio bias within species and between species is scarce. How differentiated are sex ratios among diverging species, especially when they adapt to new environments and adopt new life histories? Is sex ratio bias fixed or polymorphic within species? To answer these questions, one needs to survey sex ratios for multiple individuals per species across a lineage.

The genus *Caenorhabditis* provides a good opportunity to study the evolution of sex ratios as it is a species-rich genus [26], comprising ecologically diverse species with various life histories, population structures, and even reproductive modes [27]. Most *Caenorhabditis* species have standard female-male reproduction (dioecy), however, three species, *C. elegans, C. briggsae*, and *C. tropicalis*, have independently evolved reproduction through hermaphrodites and facultative males (androdioecy) [28, 29]. Species of both reproductive modes in *Caenorhabditis* have the same chromosomal sex determination system with females (or hermaphrodites) being XX and males being XO [30]. But unlike the male-female species that have obligate out-crossing, the androdioecious species have two ways of reproduction: a hermaphrodite can either self or out-cross with a male. Self-fertilization produces mostly hermaphrodites while males are produced by spontaneous nondisjunction of the X chromosome during meiosis at very low rates [31]. On the other hand, outcrossing with a male should produce hermaphrodites and males in a 1:1 ratio, according to Mendel’s first rule. In the common lab model, *C. elegans*, progenies derived from outcrossing display an equal sex ratio (while selfing produces >99.5% hermaphrodites) [31, 32]. In contrast, *C. briggsae*, yields a hermaphrodite-biased sex ratio upon outcrossing, putatively due to sperm competition where the male X-bearing sperm outcompete the nullo-X counterpart [33]. Knowledge of sex ratios and the underlying mechanisms in the other species of this genus is so far scarce.

In this study, we examined differences in progeny sex ratio across the genus *Caenorhabditis*. We first assayed the sex ratios for multiple strains per species for a total of 23 species, where males and females (or hermaphrodites) have continuous access to each other throughout their lifetime (“unlimited mating experiment”). As LaMunyon and Ward (1997) found sperm competition between male X-bearing and nullo-X sperm as a putative mechanism underlying the hermaphrodite-biased sex ratios in *C. briggsae* [33], we also investigated the role of sperm competition in sex ratio bias in the other *Caenorhabditis* species. In contrast to the unlimited mating experiment, we assayed the sex ratios by limiting mating to a short time interval for one strain of each species. With limited amounts of sperm, when X-bearing sperm are used up, the nullo-X sperm have a chance to catch up. Species or strains exhibiting lower female-biased sex ratios in the limited mating experiment compared to the unlimited mating experiment suggest sperm competition as a mechanism explaining sex ratio bias. These surveys and analyses would enhance our understanding of the evolution of sex ratio within this diverse genus and also shed light on the mechanisms underlying sex ratio bias.

## Results

### Sex ratio polymorphism is common in *Caenorhabditis*

When the females (or hermaphrodites) and males had continuous access to each other (unlimited mating experiment), out of a total of 106 strains (*C. briggsae* with 32 strains, 18 species with 2-5 strains, and 4 species with a single strain) tested, 72 had significant female(hermaphrodite)-biased sex ratios, 33 had non-significant bias, and only 1 strain (NIC1931, *C. imperialist* had a significant male-biased sex ratio (Table S1). For the 19 species for which we tested multiple strains, 12 showed inconsistent sex ratios between strains, i.e., some strains showed significant female-biased sex ratios (adjusted *P* <0.05, one-sided binomial test, Figure 1a) while other strains did not show sex ratio bias. Five species showed consistent female bias across strains (*C. latens*, 5 strains; *C. remanei*, 3 strains; *C*. sp. 33, 3 strains; *C*. sp. 44, 2 strains; and *C. portoensis*, 4 strains). *C. elegans* (3 strains) and *C*. sp. 41 (2 strains) had consistent non-biased sex ratios across all strains. For the 4 species for which we had only 1 strain, 2 showed significant female-biased sex ratios (*C. becei* and *C. vivipara*), whereas the other 2 did not (*C. panamensis* and *C. wallacei*).

**Figure 1.**
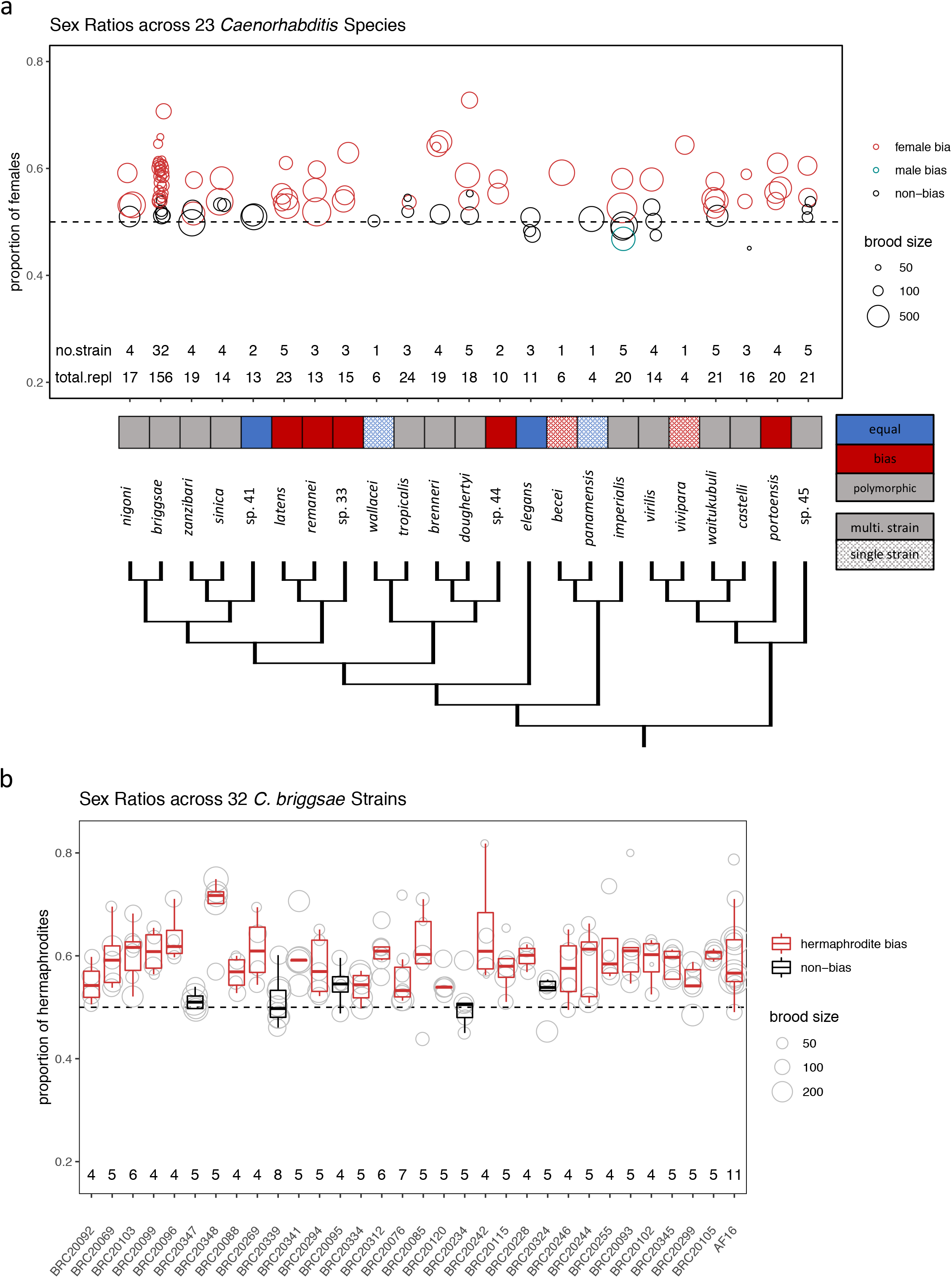
Polymorphic sex ratios across 23 *Caenorhabditis* species. (a) The proportion of females (hermaphrodites) in total progeny in the unlimited mating assay. Each circle represents a tester strain and the circle size denotes the average number of progeny across replicates. The numbers of strains (no.strain) and total replicates (total.repl) are indicated below the corresponding boxes (detailed information in Table S1). The circle color indicates whether the strain exhibited female bias, no bias, or male bias (adjusted *P* < 0.05, binomial test). The cells underneath the circle plot indicate whether the species have consistent female bias (red), consistent no bias (blue), or polymorphic sex ratios (grey) across strains. Cells with solid fill indicate species with multiple strains whereas patterned fill indicate species with a single strain. (b) Sex ratios across 32 *C. briggsae* strains. Each circle represents a replicate mating pair and the circle size denotes the total number of progeny of the replicate. The numbers of replicates are indicated below the box plots (detailed information in Table S1). The color of each box indicates whether or not the sex ratio was hermaphrodite-biased (adjusted *P* < 0.05, binomial test).

Among the 5 species with consistent female bias, 3 (*C. latens, C. remanei*, and *C*. sp. 33) form a monophyletic clade in the *Elegans* group (Figure 1a). The other 2, *C*. sp. 44 and *C. portoensis*, were in another lineage of the *Elegans* group and in the *Drosophilae* supergroup, respectively. The 2 species with consistent non-bias (*C. elegans* and *C*. sp. 41) were distributed in different linages of the *Elegans* group. The remaining 12 species with polymorphic sex ratios were scattered across the genus phylogeny, i.e., across the *Elegans* group, the *Japonica* group, and the *Drosophilae* supergroups.

Three androdioecious species have independently evolved in the *Elegans* group [29] and their sex ratio patterns differed amongst each other. *C. elegans* had consistent non-bias across strains (adjusted *P* > 0.05 for all 3 strains, one-sided binomial test, Benjamini-Hochberg correction, same for below). For *C. tropicalis*, 1 strain (BRC20400) had significant hermaphrodite bias (adjusted *P* = 0.0025) whereas the other 2 strains did not have significant bias (adjusted *P* > 0.05). For *C. briggsae*, which we had extensive sampling in Taiwan, we surveyed 31 strains in addition to the common lab strain AF16. Twenty-six strains showed significant hermaphrodite-biased sex ratios (adjusted *P* <0.05, Figure 1b, Table S1), consistent with AF16. Thus, although sex ratio bias is polymorphic across *C. briggsae* strains, the hermaphrodite-biased sex ratios dominate.

### Inter-strain crosses yielded consistent sex ratio bias

To test the possibility that sex ratio bias was due to sex-specific consequences of potential inbreeding depression [34], for 4 female-male species (*C. nigoni, C. latens, C. remanei*, and *C*. sp. 33), we performed inter-stain crosses. The strains we used for inter-strain crosses were all female-biased from the intra-strain crosses we mention above. The interstrain crosses yielded consistent female-biased sex ratios as the intra-strain crosses of the parental stains (adjusted *P* < 0.05, Table S2). These results suggest the polymorphic sex ratios we detected among species were not likely due to sex-specific consequences of inbreeding depression.

### Analyses of variation revealed significant differences between strains and species

Our results suggest that sex ratio varies both between and within strains and species. To test whether sex ratios are differentiated by strains and/or by species, we conducted a two-factor ANOVA test, which included sex ratio data of all replicates from the 106 total strains nested within 23 species. The grand ANOVA model revealed that both species and strain have significant effects on sex ratio (*P* = 4.58e-12 for species and *P* =5.79e-14 for strain, ANOVA). This suggests heritability and genetic differentiation to a large extent underlying the sex ratio profiling, despite the great variation within strains and polymorphism within species.

### Sex ratios in the limited mating experiment

To interrogate the role of sperm competition in progeny sex ratio bias, we conducted limited mating experiments to assay sex ratios for 1 strain per species for all the 23 species tested above. Our working hypothesis is that, if X-bearing sperm out-compete nullo-X sperm, we will find a female-biased sex ratio; but if sperm are limited, all sperm will have a chance to fertilize despite the competition between sperm, leading to an equal sex ratio. Species or strains exhibiting female-biased sex ratios in the unlimited mating experiment but no bias in the limited mating experiment suggest sperm competition. The 23 retested strains include 8 that showed no bias in the unlimited mating experiment and 15 that showed female-biased sex ratios. For these 23 strains, when exposed to males for only up to 5 hours, the females (or hermaphrodites) sired on average 51% (range 20-80%) fewer progeny (outcrossed progeny for androdioecious species) compared to unlimited mating, except for *C. sinica* and *C. latens*, which had comparable numbers of progeny between the two mating experiments. Out of the 8 strains that did not show female bias in the unlimited mating experiment, BRC20276 of *C*. sp. 41 showed a significantly female-biased sex ratio in the limited mating experiment (adjusted *P* = 0.0002). The remaining 7 strains consistently showed no bias in both the unlimited and limited mating experiments.

All 15 strains that showed female(hermaphrodite)-biased sex ratios in the unlimited mating experiment, also showed female(hermaphrodite)-biased sex ratios in the limited mating experiment (adjusted *P* < 0.05, binomial test; Figure 2a, Table S3). When comparing the sex ratios between unlimited and limited mating for the same strain, QG704 of *C. becei* showed significantly lower female bias under limited mating (adjusted *P* < 0.05, t-test), though the sex ratios were significantly female-biased in both experiments. This result would be compatible with sperm competition, which is either incomplete, acting along with additional female biasing processes, or both. The other 14 strains did not have significant differences between the two experiments (Table S4), possibly indicating the lack of sperm competition in these species.

**Figure 2.**
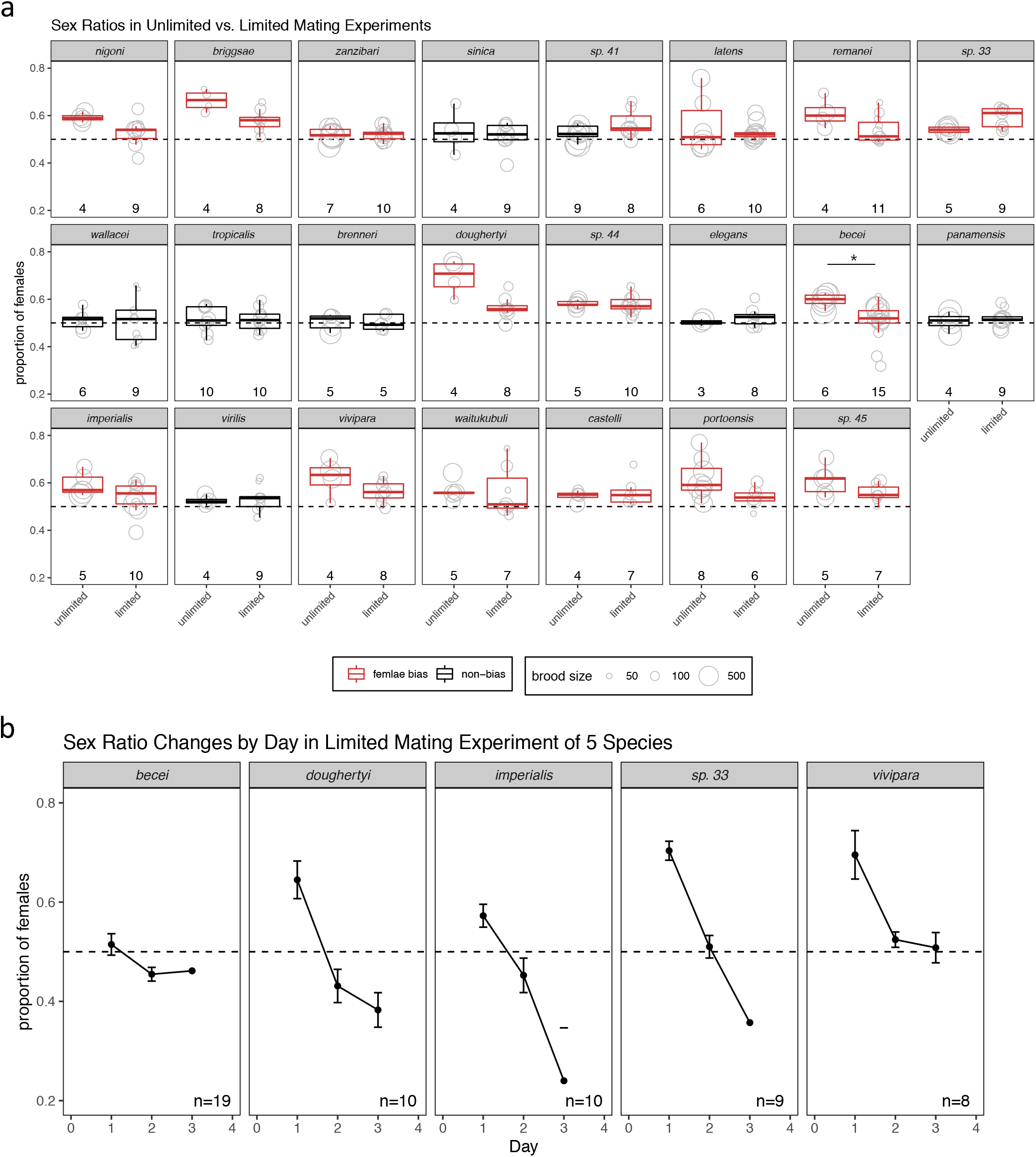
Sex ratios in the limited mating assay. (a) Sex ratios under unlimited vs. limited mating for the same strain for each of the 23 species. Each circle represents a replicate mating pair of the tester strain and the circle size denotes the total number of progeny of the replicate, with the numbers of replicates indicated below the boxplot (detailed information in Table S3). The color of each box indicates whether or not the sex ratio exhibited female bias (adjusted *P* < 0.05, binomial test). The asterisk indicates significant difference between sex ratios between the two mating experiments (adjusted *P* < 0.05, t-test). (b) Day-by-day sex ratios under limited mating for five strains (species) with linear declines in female bias over time (adjusted *P* < 0.05, linear mixed effect model). The points represent mean sex ratios per day across replicates. Error bars represent standard error of the mean. The numbers of replicates are indicated (detailed information in Table S7). The day-by-day sex ratios calculated from less than 10 progeny were excluded.

We looked for sperm competition in a second way. Although we did not observe equal sex ratios based on the total brood, the depletion of the X-bearing sperm under the limited mating condition would result in more nullo-X sperm fertilizing the oocytes over time. Therefore there should be a change in sex ratio over time. When we examined sex ratio changes against time for each strain tested under limited mating, we found heterogeneous profiles across the species (Figure S1). We identified 5 strains with linear decreases of female bias by time (BRC20005 of *C. sp. 33*, JU1333 of *C. doughertyi*, QG704 of *C. becei*, JU1905 of *C. imperialis*, and NIC1070 of *C. vivipara;* adjusted *P* < 0.05, linear mixed effect models; Figure 2b, Table S5). For these 5 strains the declines in female proportions by time suggest sperm competition as a mechanism contributing to the female bias, although other processes are likely involved because equal sex ratios were not reached for the total brood. Furthermore, for the other 10 strains without detectible sperm competition, our results imply that other processes are acting to produce female-biased sex ratios.

## Discussion

We found prevalent polymorphism in sex ratios among the *Caenorhabditis* species. The majority of strains showing sex ratio bias was female biased. Twelve out of 19 species showed polymorphic sex ratios across strains, whereas only 7 species showed consistent bias or non-bias across strains. We also compared two experimental strategies, unlimited mating versus limited mating, to investigate the contribution of sperm competition in the biased sex ratios. Most retested strains (22 out of 23) showed consistent sex ratio bias or no bias between the two experiments, suggesting that sperm competition alone cannot explain the female bias we detected in the genus. Only 1 strain had significantly lower female-biased sex ratios in the limited mating experiment than in the unlimited experiment, and 5 strains showed linear declines in female proportions along time, which are consistent with sperm competition. Our genus-wide survey of progeny sex ratios provides rich information on the evolution of sex ratios in this genus, and suggest other factors than sperm competition contribute to the sex ratio bias.

For dioecious species where sex is separated into males and females, Fisher’s principle proposes equal sex ratios as the optimum at the panmixia population level (i.e., all individuals could mate with anyone else), whereas LMC and other theories predict biased sex ratios being favored for individuals with constrained local structures. Our strategy of conducting mating crosses within isogenic strains was to assay progeny sex ratios at individual (i.e., strain) levels, whereas the sex ratios at the population level in the wild are still elusive. Nevertheless, we found many strains of the female-male species with a female-biased sex ratio in the progeny, suggesting LMC or similar forces act on these strains. *Caenorhabditis* species dwell in ecologically diverse habitats, ranging from cattle auditory canals to rotting fruits and man-made compost [27]. Despite the diverse habitats, many of these female-male *Caenorhabditis* species probably share some features of life history with their androdioecious relative *C. elegans*, such as boom-and-bust population growth in ephemeral habitats, active dispersal seeking, and strong founder effect followed by population re-expansion [35–37]. These life history features may result in high inbreeding rates and intense local competition for mating between kin. Hence, a female-biased sex ratio may be favored, according to the theory of LMC. The polymorphic sex ratios we found in many species suggest varying degrees of LMC that the different strains experience, possibly reflecting heterogeneous life histories and dispersal constraints for strains within species and/or between species. So far, for female-male species in *Caenorhabditis*, except for a few species that have been intensively sampled, such as *C remanei* [35] and *C. inopinata* [38], most other species have been sampled very rarely and thus their natural habitats are largely unknown. More knowledge about the ecology and natural history of these female-male *Caenorhabditis* species would help to explain the driving forces of sex ratio evolution in these species.

We also included the 3 androdiecious species in our sex ratio survey. *C. elegans* showed consistent no bias, *C. tropicalis* showed polymorphic sex ratio bias across three strains, whereas *C. briggsae*, where we had extensive sampling, also showed polymorphic sex ratios but hermaphrodite-biased sex ratio dominated. Again, these sex ratios of progeny we surveyed are sex ratios at the individual level. As the hermaphrodites can self-fertilize, the individual-level sex ratios can be largely decoupled from the population level. Male frequencies in natural populations have been found to be very low and out-crossing was estimated to be rare [39–41]. Though outcrossing could facilitate rapid adaptive evolution when confronted with pathogen [42], these self-fertilizing species show degraded male functions [43], such as reduced mating success [44, 45] and sperm competence [46, 47] in males of *C. elegans* and *C. briggsae*. The hermaphrodite bias we found from the mating tests in *C. briggsae* and in *C. tropicalis* support the possibility that reduced production of males in these androdiecious species might be favored.

Regardless of the reproductive mode, from a phylogenic point of view, the polymorphic sex ratios we detected in 12 out of the 19 species might be a trans-species polymorphism in the genus. Trans-species polymorphisms are genetic variation from common ancestors that are maintained through speciation. Empirical data for trans-species polymorphism have so far been limited, and mainly focus on immune-related functions, e.g., major histocompatibility complex in vertebrates [48, 49]. Trans-species polymorphism of sex ratio bias in *Caenorhabditis* nematodes would suggest strong balancing selection on sex ratios, probably due to varying intensities of LMC. Alternatively, the sex ratio bias might have evolved repeatedly within each species. Experimental evolution showed that the spider mite *Tetranychus urticae* evolved female-biased sex ratios under high LMC intensity after 54 generations, suggesting sex ratio bias could be fast evolving [50]. The possible frequent gain or loss of sex ratio bias among our *Caenorhabditis* species would reflect the adaptation to the respective habitats and life histories in these species.

Sex ratio bias under LMC has been mostly investigated in haplodiploid organisms, such as fig wasps, parasitic wasps, and mites, where sex ratios might be subject to plastic or facultative adjustments, presumably because females can control if an egg is fertilized [51]. In contrast, the nematodes we study are diploid and use chromosomal sex determination. We speculate that the polymorphic sex ratio bias is at least partially governed by genetic differences. Consistently, the ANOVA model indicates significant differences in sex ratios between strains and between species, which suggests genetic differences contribute to variation in sex ratios between strains and species.

To investigate sperm competition as a potential mechanism underlying the sex ratio bias, as previously found in *C. briggsae* [33], we conducted mating experiments where mating was limited for a few hours as opposed to the unlimited mating experiments where mating was allowed for the entirety of adulthood. A contrast of no bias in the limited mating experiment versus female bias in the unlimited mating experiment would suggest sperm competition. In the unlimited mating experiments, the couples probably mated repeatedly and the X-bearing sperm would be refilled and therefore X-bearing sperm would always take precedence over the nullo-X sperm, resulting in an overall female-biased sex ratio. In contrast, in the limited mating experiments, the amount of sperm transferred was restricted and all sperm were presumably used, resulting in an overall equal sex ratio. Surprisingly, all the 15 strains that showed significant female-biased sex ratios in unlimited mating also showed female bias in limited mating, suggesting that sperm competition alone could not explain the female bias we found in these strains. We found some hints of sperm competition in 5 strains that showed declining female bias by day under limited mating, and 1 (QG704 of *C. becei*) of them showed significantly reduced female bias in limited mating experiment compared to the unlimited mating experiment. In *C. briggsae*, LaMunyon and Ward (1997) conducted 3-hour mating as well as 8-hour mating experiments and found the overall sex ratio was equal after 3-hour mating whereas it was hermaphrodite-biased after 8-hour mating [33]. Here, we found a hermaphrodite-biased sex ratio after 5 hours of mating. These observations suggest that the amount of sperm ejaculated into the hermaphrodite is a limiting factor for sperm competition and hence sex ratio bias. Taken together, our limited sperm experiments suggest that sperm competition may at least partially contribute to female-biased sex ratios in some cases, though the prevalent sex ratio bias across the genus are largely not explained by sperm competition.

The presence of selfish genetic elements could provide an alternative mechanism for sex ratio bias where the proportion of X-bearing sperm is selfishly enhanced, such as *segregation distorter* on the X chromosome of *Drosophila simulans* [52]. So far, several selfish elements have been discovered in *Caenorhabditis* nematodes, but they all reside on autosomes [53–56]. Other potential mechanisms include non-canonical meiosis, such as asymmetric division in gametogenesis in *Auanema* (aka *Rhabditis* sp.) nematodes, where XO males exclusively produce X-bearing sperm and hermaphrodites produce mostly nullo-X oocytes and diplo-X sperm [57–60].

While equal sex ratios are presumably predominant in nature, biased sex ratios may be largely underappreciated. In this study, we carried out a broad survey of sex ratio bias in outcrossed progeny across *Caenorhabditis* nematodes. Our findings of prevalent sex ratio polymorphism add to the limited knowledge about sex ratio bias in the animal kingdom, and provide empirical support that sex ratio bias can evolve rapidly within a single genus.

## Materials and Methods

### Species and culture

We examined 23 *Caenorhabditis* species for progeny sex ratio in this study, including 4 new species: *C*. sp. 33, *C*. sp. 41, *C*. sp. 44, and *C*. sp. 45. These 4 new species were placed onto the phylogenetic tree based on their ITS2 sequences (the intergenic region between the 5.8S and LSU rRNA genes) [61, 62]. These 23 species comprise 17 species from the *Elegans* supergroup and 6 from the *Drosophila* supergroup. Of the 17 species from the *Elegans* supergroup, 3 were from the *Japonica* group and the rest were from the *Elegans* group, including the 3 androdioecious species (*C. elegans, C. briggsae*, and *C. tropicalis*). Except for *C. briggsae*, we included 1-5 strains for each species (4 species with only 1 strain, 2 species with 2 strains, 5 species with 3 strain, 6 species with 4 strains, and 5 species with 5 strains). For *C. briggsae*, we had 32 strains. (Table S1). All worms were grown at room temperature (23-24°C) on nematode growth media agar plates seeded with OP50 *Escherichia coli* bacteria.

### Sex ratio assay

#### Mating experimental design

For the female-male species, sex ratios were assayed by intra-strain crosses, i.e., females and males from the same wild-type isofemale strains (tester strains). For the androdioecious species, we crossed males from wild-type strains (tester strain) to hermaphrodite strains carrying a recessive mutation, so that the outcrossed progeny were visually identifiable. The recessive morphological mutant strains had Uncoordinated (Unc) or Dumpy (Dpy) phenotypes: *C. elegans* (BRC0189, *unc-119(ed9)); C. briggsae* (BRC0258, *unc(ant10));* and *C. tropicalis* (BRC0419, *dpy(ant23*)). For each of these androdioecious species, we had multiple tester strains, especially for *C. briggsae*, for which we had many isolates collected in Taiwan.

We set up mating experiments to assay sex ratio in the same manner across all the species tested. In the “unlimited mating” experiment, we placed one L4 female (or hermaphrodite) and one L4 male on a fresh 55 mm diameter Petri plate and then transferred them together every day to fresh plates until they died or produced no more eggs. When the progeny reached the L4 or adult stage, the numbers of outcrossed females (or hermaphrodites) and males per plate were manually scored under the microscope. For the male-female species, all progeny were counted, while for the androdioecious species, only wild-type cross progeny were counted whereas selfed Unc or Dpy progeny were ignored. Note, while we did not systematically survey numbers of inviable embryos or young larvae, we did not notice any appreciable amounts of dead eggs (i.e., greater than a few) that could have accounted for the reduced number of males. Mating pairs with total number of out-crossed progeny smaller than 40 were excluded from further analyses to ensure adequate statistical power. For each test of the strains, we had at least three replicate mating pairs. The counts of progeny of the two sexes for each day and for each replicate were used for further statistical analyses (see below).

In addition to intra-strain crosses, for 4 female-male species (*C. nigoni, C. latens, C. remanei*, and *C*. sp. 33), we also conducted inter-strain crosses to test the possibility that sex ratio bias was due to sex-specific consequences of inbreeding depression [34]. For *C. nigoni*, we intercrossed 2 strains (BRC20235 and BRC20079, reciprocally) and then crossed the heterozygous F1 males with the maternal strain. For the 3 species, *C. latens, C. remanei*, and *C*. sp. 33, for which we used 3 strains for inter-strain crosses, we first crossed 2 strains and then crossed the heterozygous F1 males with the third strain (Table S2). We scored the two sexes in the F2 progeny in the same manner as described above for the “unlimited mating” experiment.

#### Time limitations for mating

To interrogate the potential role of competition between male X-bearing sperm and the nullo-X counterpart in progeny sex ratio bias, we conducted sex ratio assays by “limited mating” for one tester strain per species (Table S3). To do so, L4 females (hermaphrodites) and L4 males were isolated one day prior to the cross to ensure their virginity and matured singly overnight. The next day, one male was added to one isolated female (hermaphrodite). The male was removed when mating plugs were observed on the females (hermaphrodites) or after 5 hours. Mated females (hermaphrodites) were transferred daily to new plates. For *C. tropicalis*, all pairs failed to mate within 5 hours, so we crossed one hermaphrodite with three males to increase the chance of mating. The progeny sex ratios were scored as described above.

### Statistical analysis

For each of the sex ratio assays, unlimited mating or limited mating, we tested whether the progeny sex ratio was biased. For each tester strain, the counts of total females (hermaphrodites) and total males from all days and all replicates were summed up and used for the binomial test. If the crosses yielded more female (hermaphrodite) than male progeny, we tested whether the proportions of females (hermaphrodites) significantly exceeded equality, and vice versa (R, binom.test, alternative = “greater” for more females and alternative = “less” for more males). The *P*-values of the strains were respectively corrected for multiple testing for the total numbers of strains using the Benjamini-Hochberg method [63]. Due to the unbalanced sample sizes, the 31 *C. briggsae* strains were corrected separately from the other 75 strains of other species. Strains with the corrected *P*-values smaller than 0.05 were defined as having sex ratio bias. Species with strains having sex ratio bias and also strains without bias are defined as species with polymorphic sex ratios. For the unlimited mating experiment with multiple strains, to analyse sex ratio variation within strains, between strains, and between species, we used an ANOVA model with species nested with strains (aov function from R package lme4, female_proportion ~ species/strain). To compare the sex ratios between unlimited mating and limited mating for the same strains, we tested the differences in sex ratios between these two experiments by a t-test. To estimate the sex ratio changes by day in the limited mating experiment, we built linear mixed models including day as fixed effects and replicates as random effects for each strain (lme function from R package nlme, female_proportion ~ day, random=~1|replicate). All statistical analyses were performed in R v.3.2.3 except otherwise indicated [64].

## Acknowledgements

Some *Caenorhabditis* wild isolates were kindly provided by Marie-Anne Félix and Karin Kiontke, and additional strains were provided by the *C. elegans* Genetic Center (CGC), which is funded by the National Institutes of Health Office of Research Infrastructure Programs (P40 OD010440). We thank Matthew Rockman for sharing and discussing unpublished results with us and Marie-Anne Félix for her comments and suggestions on the manuscript.

## Author contributions

Y.-H.L., and J.W. conceived and designed the study. Y.-H.L, Y.H., J.-C.H., T.S.L, F.-J.Y., T.C., and J.W. conducted experiments. C.B. contributed strains and help revised the manuscript. Y.H. and Y.-H.L. analyzed the data. Y.H., Y.-H.L., and J.W wrote the manuscript with input from others.

## Ethics statement

This research was exempt from ethics approvals for the animal treatments because the studied species are not considered under the animal treatment guidelines.

## Data Accessibility

The datasets supporting this article have been uploaded as part of the supplementary material.

## Supplementary Figures and Tables

Figure S1. Day-by-day sex ratios of strains tested for unlimited and limited mating experiments, the x-axis represents days after mating.

Table S1. Sex ratios of all 106 strains tested in the unlimited mating experiment. Female and male counts were pooled from all replicates. These are the data used for Figure 1.

Table S2. Sex ratios of inter-stain crosses

Table S3. Sex ratios of strains tested in the limited mating experiment. Female and male counts were pooled from all replicates. These are the data used for Figure 2a.

Table S4. Results of t-tests between unlimited mating and limited mating for 23 retested strains

Table S5. Results of linear mixed effect models of sex ratios against time in the limited mating experiment

Table S6. Sex ratio count data by day for unlimited mating experiment from all replicates of strains.

Table S7. Sex ratio count data by day for limited mating experiment from all replicates of strains. These are the data used for Figure 2b.

## References

[1] Darwin, C. 1874 The descent of man and selection in relation to sex. 2nd ed, John Murray, London.

[2] Edwards, A.W.F. 1998 Natural selection and the sex ratio: Fisher’s sources. The American Naturalist 151, 564–569. (doi:10.1086/286141).

[3] Fisher, R.A. 1930 The genetical theory of natural selection, Clarendon, Oxford.

[4] Hamilton, W.D. 1967 Extraordinary sex ratios. Science 156, 477–488.

[5] West, S.A., Herre, E.A. & Sheldon, B.C. 2000 The benefits of allocating sex. Science 290, 288–290. (doi:doi:10.1126/science.290.5490.288).

[6] Charnov, E.L. 1982 The theory of sex allocation, Princeton University Press, Princeton, NJ.

[7] Clark, A.B. 1978 Sex ratio and local resource competition in a prosimian primate. Science 201, 163–165. (doi:10.1126/science.201.4351.163).

[8] Gowaty, P.A. & Lennartz, M.R. 1985 Sex ratios of nestling and fledgling red-cockaded woodpeckers (*Picoides borealis*) favor males. The American Naturalist 126, 347–353. (doi:10.1086/284421).

[9] Emlen, S.T., Emlen, J.M. & Levin, S.A. 1986 Sex-ratio selection in species with helpers-at-the-nest. The American Naturalist 127, 1–8. (doi:10.1086/284463).

[10] Trivers, R.L. & Willard, D.E. 1973 Natural selection of parental ability to vary the sex ratio of offspring. Science 179, 90–92. (doi:10.1126/science.179.4068.90).

[11] Paul, R.E., Raibaud, A. & Brey, P.T. 1999 Sex ratio adjustment in *Plasmodium gallinaceum*. Parassitologia 41, 153–158.

[12] Osgood, S.M., Eisen, R.J. & Schall, J.J. 2002 Gametocyte sex ratio of a malaria parasite: experimental test of heritability. The Journal of parasitology 88, 494–498. (doi:10.1645/0022-3395(2002)088[0494:Gsroam]2.0.Co;2).

[13] Reece, S.E., Drew, D.R. & Gardner, A. 2008 Sex ratio adjustment and kin discrimination in malaria parasites. Nature 453, 609–614. (doi:10.1038/nature06954).

[14] Ferguson, D.J. 2002 *Toxoplasma gondii* and sex: essential or optional extra? Trends in parasitology 18, 355–359.

[15] Haukisalmi, V., Henttonen, H. & Vikman, P. 1996 Variability of sex ratio, mating probability and egg production in an intestinal nematode in its fluctuating host population. International Journal for Parasitology 26, 755–764. (doi:https://doi.org/10.1016/0020-7519(96)00058-6).

[16] Kloch, A., Michalski, A., Bajer, A. & Behnke, J. 2015 Biased sex ratio among worms of the family *Heligmosomidae--searching* for a mechanism. Int J Parasitol 45, 939–945. (doi:10.1016/j.ijpara.2015.08.002).

[17] Herre, E.A. 1985 Sex ratio adjustment in fig wasps. Science 228, 896–898. (doi:10.1126/science.228.4701.896).

[18] Werren, J.H. 1980 Sex ratio adaptations to local mate competition in a parasitic wasp. Science 208, 1157–1159. (doi:doi:10.1126/science.208.4448.1157).

[19] David M. Shuker, Ido Pen, Alison B. Duncan, Sarah E. Reece & Stuart A. West. 2005 Sex ratios under asymmetrical local mate competition: theory and a test with parasitoid wasps. The American Naturalist 166, 301–316. (doi:10.1086/432562).

[20] Griffiths, N.T. & Godfray, H.C.J. 1988 Local mate competition, sex ratio and clutch size in *bethylid* wasps. Behavioral Ecology and Sociobiology 22, 211–217.(doi:10.1007/BF00300571).

[21] Komdeur, J. 1996 Facultative sex ratio bias in the offspring of Seychelles warblers. Proceedings of the Royal Society B: Biological Sciences 263, 661–666. (doi:doi:10.1098/rspb.1996.0099).

[22] Komdeur, J., Daan, S., Tinbergen, J. & Mateman, C. 1997 Extreme adaptive modification in sex ratio of the Seychelles warbler’s eggs. Nature 385, 522–525. (doi:10.1038/385522a0).

[23] Malcolm, J.R. & Marten, K. 1982 Natural selection and the communal rearing of pups in African wild dogs (Lycaon pictus). Behavioral Ecology and Sociobiology 10, 1–13.(doi:10.1007/BF00296390).

[24] Clutton-Brock, T.H., Albon, S.D. & Guinness, F.E. 1984 Maternal dominance, breeding success and birth sex ratios in red deer. Nature 308, 358–360. (doi:10.1038/308358a0).

[25] Symington, M.M. 1987 Sex ratio and maternal rank in wild spider monkeys: when daughters disperse. Behavioral Ecology and Sociobiology 20, 421–425. (doi:10.1007/BF00302985).

[26] Kiontke, K. & Fitch, D.H. 2005 The phylogenetic relationships of *Caenorhabditis* and other *rhabditids*. WormBook: the online review of C. elegans biology, 1–11. (doi:10.1895/wormbook.1.11.1).

[27] Kiontke, K. & Sudhaus, W. 2006 Ecology of *Caenorhabditis* species. WormBook: the online review of C. elegans biology, 1–14. (doi:10.1895/wormbook.1.37.1).

[28] Cutter, A.D. & Payseur, B.A. 2003 Rates of deleterious mutation and the evolution of sex in *Caenorhabditis*. Journal of evolutionary biology 16, 812–822. (doi:10.1046/j.1420-9101.2003.00596.x).

[29] Kiontke, K., Gavin, N.P., Raynes, Y., Roehrig, C., Piano, F. & Fitch, D.H.A. 2004 *Caenorhabditis* phylogeny predicts convergence of hermaphroditism and extensive intron loss. Proceedings of the National Academy of Sciences 101, 9003–9008. (doi:10.1073/pnas.0403094101).

[30] Haag, E.S. 2005 The evolution of nematode sex determination: *C. elegans* as a reference point for comparative biology. WormBook: the online review of C. elegans biology, 1–14. (doi:10.1895/wormbook.1.120.1).

[31] Hodgkin, J., Horvitz, H.R. & Brenner, S. 1979 Nondisjunction mutants of the nematode *Caenorhabditis elegans*. Genetics 91, 67–94.

[32] Teotónio, H., Manoel, D. & Phillips, P.C. 2006 Genetic variation for outcrossing among *Caenorhabditis elegans* isolates. Evolution 60, 1300–1305. (doi:https://doi.org/10.1111/j.0014-3820.2006.tb01207.x).

[33] LaMunyon, C.W. & Ward, S. 1997 Increased competitiveness of nematode sperm bearing the male X□ chromosome. Proceedings of the National Academy of Sciences 94, 185–189. (doi:10.1073/pnas.94.1.185).

[34] Ebel, E.R. & Phillips, P.C. 2016 Intrinsic differences between males and females determine sex-specific consequences of inbreeding. BMC Evol Biol 16, 36. (doi:10.1186/s12862-016-0604-5).

[35] Petersen, C., Dirksen, P., Prahl, S., Strathmann, E.A. & Schulenburg, H. 2014 The prevalence of *Caenorhabditis elegans* across 1.5 years in selected North German locations: the importance of substrate type, abiotic parameters, and Caenorhabditis competitors. BMC ecology 14, 4. (doi:10.1186/1472-6785-14-4).

[36] Félix, M.-A. & Braendle, C. 2010 The natural history of *Caenorhabditis elegans*. Current Biology 20, R965–R969. (doi:https://doi.org/10.1016/j.cub.2010.09.050).

[37] Frézal, L. & Félix, M.A. 2015 *C. elegans* outside the Petri dish. eLife 4. (doi:10.7554/eLife.05849).

[38] Kanzaki, N., Tsai, I.J., Tanaka, R., Hunt, V.L., Liu, D., Tsuyama, K., Maeda, Y., Namai, S., Kumagai, R., Tracey, A., et al. 2018 Biology and genome of a newly discovered sibling species of *Caenorhabditis elegans*. Nature Communications 9, 3216. (doi:10.1038/s41467-018-05712-5).

[39] Barrière, A. & Félix, M.A. 2005 High local genetic diversity and low outcrossing rate in *Caenorhabditis elegans* natural populations. Current biology: CB 15, 1176–1184. (doi:10.1016/j.cub.2005.06.022).

[40] Barrière, A. & Félix, M.A. 2007 Temporal dynamics and linkage disequilibrium in natural *Caenorhabditis elegans* populations. Genetics 176, 999–1011. (doi:10.1534/genetics.106.067223).

[41] Gimond, C., Jovelin, R., Han, S., Ferrari, C., Cutter, A.D. & Braendle, C. 2013 Outbreeding depression with low genetic variation in selfing *Caenorhabditis* nematodes. Evolution 67, 3087–3101. (doi:10.1111/evo.12203).

[42] Morran, L.T., Schmidt, O.G., Gelarden, I.A., Parrish, R.C., 2nd & Lively, C.M. 2011 Running with the Red Queen: host-parasite coevolution selects for biparental sex. Science 333, 216–218. (doi:10.1126/science.1206360).

[43] Cutter, A.D., Morran, L.T. & Phillips, P.C. 2019 Males, Outcrossing, and Sexual Selection in *Caenorhabditis* Nematodes. Genetics 213, 27–57. (doi:10.1534/genetics.119.300244).

[44] Garcia, L.R., LeBoeuf, B. & Koo, P. 2007 Diversity in mating behavior of hermaphroditic and male–female *Caenorhabditis* nematodes. Genetics 175, 1761–1771. (doi:10.1534/genetics.106.068304).

[45] Kleemann, G.A. & Basolo, A.L. 2007 Facultative decrease in mating resistance in hermaphroditic *Caenorhabditis elegans* with self-sperm depletion. Animal Behaviour 74, 1339–1347. (doi:https://doi.org/10.1016/j.anbehav.2007.02.031).

[46] Yin, D., Schwarz, E.M., Thomas, C.G., Felde, R.L., Korf, I.F., Cutter, A.D., Schartner, C.M., Ralston, E.J., Meyer, B.J. & Haag, E.S. 2018 Rapid genome shrinkage in a self-fertile nematode reveals sperm competition proteins. Science 359, 55–61.(doi:doi:10.1126/science.aao0827).

[47] Yin, D. & Haag, E.S. 2019 Evolution of sex ratio through gene loss. Proceedings of the National Academy of Sciences 116, 12919–12924. (doi:10.1073/pnas.1903925116).

[48] Azevedo, L., Serrano, C., Amorim, A. & Cooper, D.N. 2015 Trans-species polymorphism in humans and the great apes is generally maintained by balancing selection that modulates the host immune response. Human Genomics 9, 21. (doi:10.1186/s40246-015-0043-1).

[49] Těšický, M. & Vinkler, M. 2015 Trans-species polymorphism in immune genes: general pattern or mhc-restricted phenomenon? Journal of Immunology Research 2015, 838035. (doi:10.1155/2015/838035).

[50] Macke, E., Magalhães, S., Bach, F. & Olivieri, I. 2011 Experimental evolution of reduced sex ratio adjustment under local mate competition. Science 334, 1127–1129. (doi:doi:10.1126/science.1212177).

[51] Mueller, U.G. 1991 Haplodiploidy and the evolution of facultative sex ratios in a primitively eusocial bee. Science 254, 442–444. (doi:doi:10.1126/science.254.5030.442).

[52] Tao, Y., Araripe, L., Kingan, S.B., Ke, Y., Xiao, H. & Hartl, D.L. 2007 A sex-ratio meiotic drive system in *Drosophila simulans*. ii: an x-linked distorter. PLOS Biology 5, e293. (doi:10.1371/journal.pbio.0050293).

[53] Seidel, H.S., Rockman, M.V. & Kruglyak, L. 2008 widespread genetic incompatibility in *C. elegans* maintained by balancing selection. Science 319, 589–594. (doi:10.1126/science.1151107).

[54] Ben-David, E., Burga, A. & Kruglyak, L. 2017 A maternal-effect selfish genetic element in *Caenorhabditis elegans*. Science 356, 1051–1055. (doi:10.1126/science.aan0621).

[55] Ben-David, E., Pliota, P., Widen, S.A., Koreshova, A., Lemus-Vergara, T., Verpukhovskiy, P., Mandali, S., Braendle, C., Burga, A. & Kruglyak, L. 2021 Ubiquitous selfish toxin-antidote elements in *Caenorhabditis* species. Current Biology 31, 990–1001.e1005. (doi:https://doi.org/10.1016/j.cub.2020.12.013).

[56] Noble, L.M., Yuen, J., Stevens, L., Moya, N., Persaud, R., Moscatelli, M., Jackson, J.L., Zhang, G., Chitrakar, R., Baugh, L.R., et al. 2021 Selfing is the safest sex for *Caenorhabditis tropicalis*. eLife 10, e62587. (doi:10.7554/eLife.62587).

[57] Shakes, D.C., Neva, B.J., Huynh, H., Chaudhuri, J. & Pires-daSilva, A. 2011 Asymmetric spermatocyte division as a mechanism for controlling sex ratios. Nature Communications 2, 157. (doi:10.1038/ncomms1160).

[58] Kanzaki, N., Kiontke, K., Tanaka, R., Hirooka, Y., Schwarz, A., Müller-Reichert, T., Chaudhuri, J. & Pires-daSilva, A. 2017 Description of two three-gendered nematode species in the new genus Auanema (Rhabditina) that are models for reproductive mode evolution. Scientific Reports 7, 11135. (doi:10.1038/s41598-017-09871-1).

[59] Winter, E.S., Schwarz, A., Fabig, G., Feldman, J.L., Pires-daSilva, A., Müller-Reichert, T., Sadler, P.L. & Shakes, D.C. 2017 Cytoskeletal variations in an asymmetric cell division support diversity in nematode sperm size and sex ratios. Development 144, 3253–3263. (doi:10.1242/dev.153841).

[60] Tandonnet, S., Farrell, M.C., Koutsovoulos, G.D., Blaxter, M.L., Parihar, M., Sadler, P.L., Shakes, D.C. & Pires-daSilva, A. 2018 Sex-and gamete-specific patterns of X chromosome segregation in a trioecious nematode. Current Biology 28, 93–99.e93. (doi:https://doi.org/10.1016/j.cub.2017.11.037).

[61] Kiontke, K.C., Félix, M.-A., Ailion, M., Rockman, M.V., Braendle, C., Pénigault, J.-B. & Fitch, D.H.A. 2011 A phylogeny and molecular barcodes for *Caenorhabditis*, with numerous new species from rotting fruits. BMC Evolutionary Biology 11, 339. (doi:10.1186/1471-2148-11-339).

[62] Félix, M.-A., Braendle, C. & Cutter, A.D. 2014 A streamlined system for species diagnosis in *Caenorhabditis* (nematoda: *rhabditidae*) with name designations for 15 distinct biological species. PLOS ONE 9, e94723. (doi:10.1371/journal.pone.0094723).

[63] Benjamini, Y. & Hochberg, Y. 1995 Controlling the false discovery rate: a practical and powerful approach to multiple testing. Journal of the Royal Statistical Society. Series B (Methodological) 57, 289–300.

[64] R Development Core Team. 2020 A language and environment for statistical computing. (Vienna, Austria, R Foundation for Statistical Computing.

