## Supplementary figures and images for "Widespread sex ratio polymorphism in *Caenorhabditis* nematodes"

### figureS1.byday_sexratio_byspecies.pdf

Sex Ratio Changes by Day in unLimited Mating

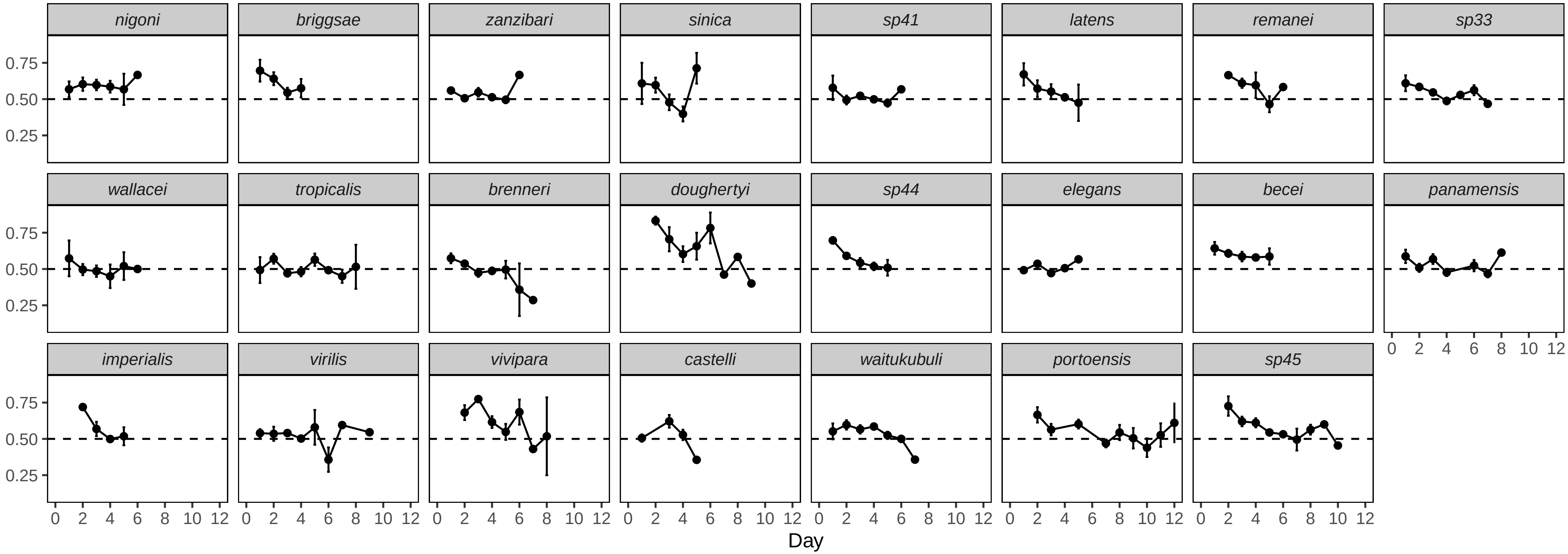


Sex Ratio Changes by Day in Limited Mating

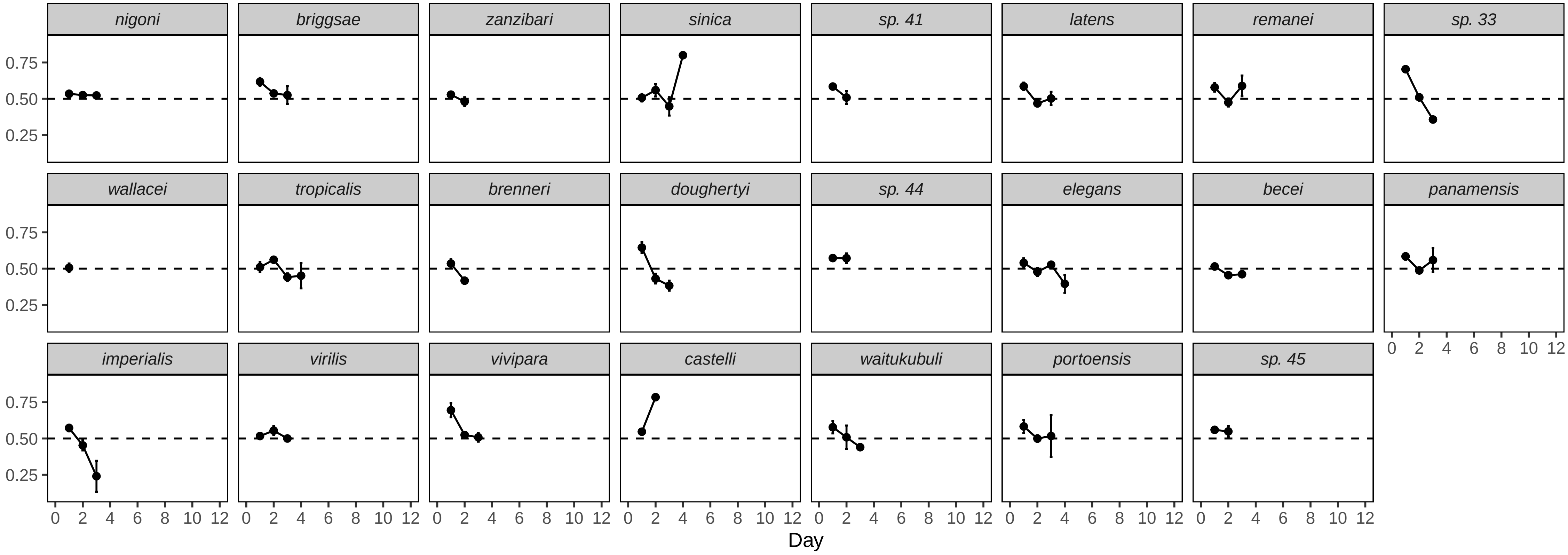
